# Associations between methylation age and brain age in late adolescence

**DOI:** 10.1101/2022.09.08.506972

**Authors:** Faye Sanders, Vilte Baltramonaityte, Gary Donohoe, Neil M Davies, Erin C. Dunn, Charlotte A.M. Cecil, Esther Walton

**Affiliations:** Department of Psychology, University of Bath, Bath, United Kingdom; School of Psychology, National University of Ireland Galway, Ireland; Centre for Neuroimaging, Cognition & Genomics, National University of Ireland, Galway, Ireland; Medical Research Council Integrative Epidemiology Unit at the University of Bristol, BS8 2BN, United Kingdom; Population Health Sciences, Bristol Medical School, University of Bristol, Barley House, Oakfield Grove, Bristol, BS8 2BN, United Kingdom; K.G. Jebsen Center for Genetic Epidemiology, Department of Public Health and Nursing, Norwegian University of Science and Technology, Norway; Psychiatric and Neurodevelopmental Genetics Unit, Centre for Genomic Medicine, Massachusetts General Hospital, Boston, MA, 02114, USA; Department of Psychiatry, Harvard Medical School, Boston, MA, 02115, USA; Stanley Center for Psychiatric Research, The Broad Institute of Harvard and MIT, Cambridge, MA, 02142, USA; Center on the Developing Child at Harvard University, Cambridge, MA, 02138, USA; Department of Child and Adolescent Psychiatry/Psychology, Erasmus MC, University Medical Center Rotterdam, Rotterdam, the Netherlands; Department of Epidemiology, Erasmus MC, University Medical Center Rotterdam, Rotterdam, the Netherlands; Molecular Epidemiology, Department of Biomedical Data Sciences, Leiden University Medical Center, Leiden, the Netherlands

**Keywords:** epigenetic age, brain age, adolescence, ALSPAC, development, DNA methylation

## Abstract

Recent research suggests that biological age, based on DNA methylation or neuroimaging measures, may predict health traits in adulthood more accurately than chronological age. However, whether these findings apply to earlier stages in life is unknown. We therefore aimed to characterise the performance of and interdependence between measures of biological age during adolescence, leveraging longitudinal data from a subsample of young adolescents from the population-based ALSPAC cohort (n=386).

We derived four methylation age measures in late adolescence (17-19 years) and a measure of brain age derived from structural neuroimaging data (18-24 years). We then examined associations between these measures of biological age, and their relationship with five measures of physical, cognitive and mental health (8-18 years).

Brain age was largely independent of different measures of methylation age, after accounting for age, cell type composition, array and study (beta range: -0.60 to 0.60, all p>0.05). Smoking and BMI were associated with three measures of advanced methylation age (beta range: -0.35 to 0.52, all p<0.05), but not brain age. Depressive symptoms and cognitive ability were unrelated to all measures of biological age.

Our findings emphasize the variability of and independence between these methylation- and brain-based measures of age in adolescents. They also highlight the importance of tracking the mosaic of ageing in younger populations.

## Introduction

Longer life expectancy and the associated increases in age-related illnesses are placing growing pressures on society. Although the risk of developing physical or mental illness increases generally as we age, there is a large amount of variability within individuals of the same chronological age. Recent research suggests that defining age biologically, rather than chronologically, may provide a more accurate predictor of health outcomes. Measures of biological age can be derived from various biological data, including epigenetic-based DNA methylation signatures and structural brain scans to characterize “brain age” (Han et al., 2021; Horvath, 2013). Advanced biological age has been associated with several health outcomes, such as cognitive decline, health span, and all-cause mortality (Belsky et al., 2015; Chen et al., 2016; Levine et al., 2018). Therefore, analytic approaches that incorporate measures of biological age, as substitutes or in addition to chronological age alone, may improve our ability to predict mortality and age-related diseases (Levine, 2013).

A growing body of research focuses on estimates of biological age based on DNA methylation profiles, known as methylation age (or epigenetic ‘clocks’). Methylation profiles consistently correlate with chronological age across various tissues, cell types and species (Horvath, 2013). Methylation age has been found to be an important predictor of mainly physical, but also some mental and cognitive health traits, especially in older individuals (Field et al., 2018; Jansen et al., 2021). For example, advanced methylation was closely associated with increased BMI, frailty and all-cause mortality in adults (Levine, 2013; Ryan et al., 2020). Critically, chronological age no longer significantly predicted health outcomes in both adults and elderly individuals (Levine, 2013) when methylation age was taken into account. Thus, these findings indicate the predictive potential of methylation age for health outcomes over and above chronological age.

However, most of these methylation age measures are based on easily accessible peripheral tissues such as blood and little is known about how these peripheral age markers relate to ageing processes in the brain. There is some evidence that biological ageing is a system-wide process, associated with a decline across tissues. For example, Hillary et al. (2021) found advanced methylation age associated with decreased brain volume and increased brain white matter hyperintensities in an elderly sample of 540 participants. Gadd and colleagues (2021) reported that methylation scores for smoking correlated across blood and brain tissue, also indicating that some methylation measures follow a system-wide pattern across the blood and brain.

Conversely, there is also evidence for substantial variability in the rate of functional and structural decline across different tissues, a concept referred to as the ‘mosaic of ageing’ (Cevenini et al., 2008; Walker and Herndon, 2010). If temporal and cross-tissue variability exists in ageing, consistency between peripheral and brain-based ageing processes cannot be presumed. Clarifying the nature and direction of associations between cross-tissue measures of age is important, as it could increase our understanding of the mechanisms and interactions within the ageing process across the lifespan and within specific developmental stages.

The recent development of two new brain-based measures of age might enable researchers to address this question. First, Shireby and colleagues (2020) developed a ‘cortical clock’ - a novel methylation age measure based on post-mortem cortical tissue across the life course, which also performs well when applied to blood tissue. Second, a neuroimaging measure of brain age has been created (Han et al., 2021). This neuroimaging measure is based on structural magnetic resonance imaging (MRI) data, including grey matter volume, cortical thickness, and surface area. Akin to the role of methylation clocks in predicting health outcomes, advanced brain age has been associated with age-related illnesses such as Alzheimer’s disease and mild cognitive impairment (Franke and Gaser, 2012; Gaser et al., 2013). Strikingly, an association between brain age and illnesses previously less defined as related to chronological age (e.g., depression, schizophrenia and epilepsy) has also been observed (Han et al., 2021; Koutsouleris et al., 2014; Sone et al., 2021), suggesting that the biological process of ageing may play a greater role in the development or symptomology of these illnesses than previously considered.

To our knowledge, only a few studies tested for associations between methylation age and brain age. In a cohort of 620 older adults, Cole et al. (2018) found little evidence that methylation correlated with brain age. However, the authors did report independent associations between mortality risk and methylation age or brain age, respectively, highlighting the unique importance of both measures for healthy ageing. Similar results were reported by Zheng et al. (2022) in a sample of 326 adults. Teeuw et al. (2021) also found little evidence that two measures of methylation age associated with brain age in 172 adults with and without schizophrenia. An additional study, using genetic data, reported low genetic correlations between different measures of methylation age and brain age (Gialluisi et al., 2021). These correlations varied in both strength and direction and across age groups, suggesting there may be more complex and developmentally patterned relationships.

So far, it is unclear to which degree methylation age and brain age associate with each other in younger, population-based samples. Studies of children and adolescents have detected associations between *either* methylation *or* brain age and markers of development, behaviour or health (Huang et al., 2019; Rudolph et al., 2017). However, research to date has not yet investigated the degree to which methylation and brain age associate *with each other* in adolescents. Knowing the relationship between these markers is critical to understand the interdependence of different biological ageing processes and how to design early and effective interventions to prevent advanced ageing.

The aim of this study was threefold:

Aim 1) to estimate the associations between chronological and biological age in a population-based cohort of young adolescents;

Aim 2) to examine the associations across modalities of biological age, (i.e., measures of methylation age and brain age, which were developed in either peripheral or cortical tissues); and Aim 3) to determine the associations between these biological age measures and physical, cognitive and mental health traits.

## Methods

### Study population

Our adolescent sample was drawn the Avon Longitudinal Study of Parents and Children (ALSPAC). Pregnant women resident in Avon, UK with expected dates of delivery between 1st April 1991 to 31st December 1992 were invited to take part in the study. The initial number of pregnancies enrolled was 14,541 (for these at least one questionnaire has been returned or a “Children in Focus” clinic had been attended by July 19 1999). Of these initial pregnancies, there was a total of 14,676 foetuses, resulting in 14,062 live births and 13,988 children who were alive at 1 year of age (Boyd et al., 2013; Fraser et al., 2013; Northstone et al., 2019). Please note the study website contains details of all the data that is available through a fully searchable data dictionary and variable search tool (http://www.bristol.ac.uk/alspac/researchers/our-data/). Ethical approval for the study was obtained from the ALSPAC Ethics and Law Committee and the Local Research Ethics Committees. Consent for biological samples has been collected in accordance with the Human Tissue Act (2004).

For this study, we restricted our analyses to young adolescents with both methylation and brain imaging data, as described in the next sections. To avoid potential biases arising from population stratification (meaning differences in genetic structure) or shared family environment, only adolescents whose parents reported white ethnicity were included in the analyses sample and – in case of twins - only the first-born twin per family. Our final analytic sample consisted of n=386 participants in late adolescence (Table 1).

**Table 1.**
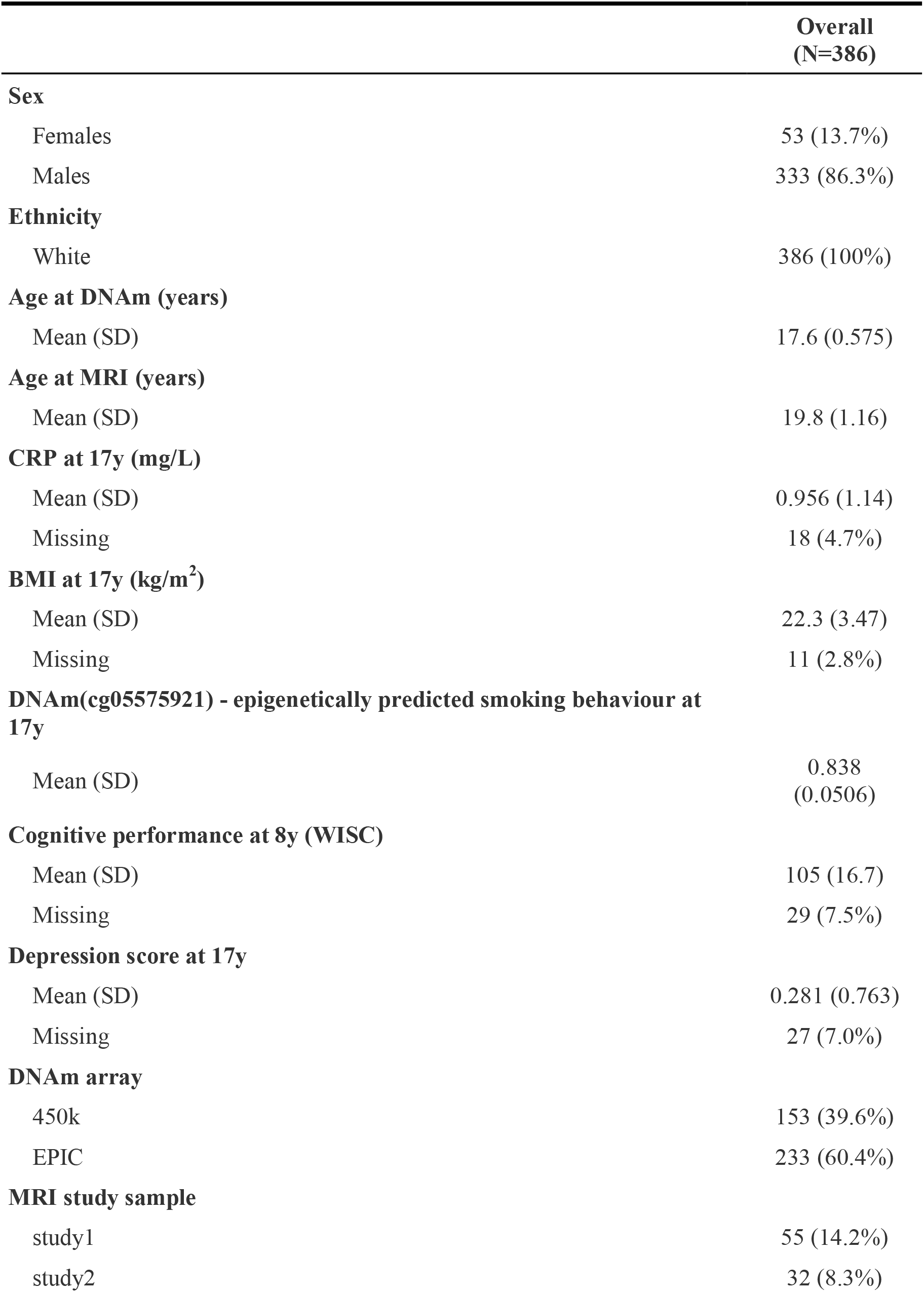
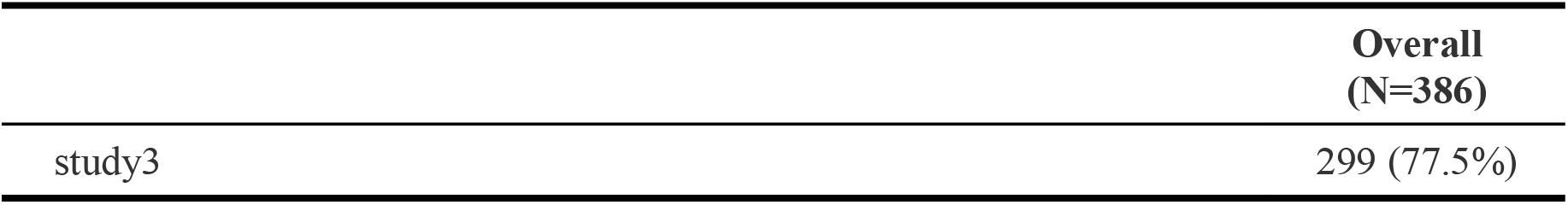
Sample demographics. DNAm = DNA methylation, MRI = magnetic resonance imaging, CRP = C reactive protein, BMI = body mass index, and WISC = Weschler Intelligence Scale for Children.

### Methylation preprocessing

Blood samples from adolescents at age 17 years were selected for analysis as part of the Accessible Resource for Integrative Epigenomic Studies (ARIES, http://www.ariesepigenomics.org.uk/; Relton et al. 2015). Bisulfite conversion was performed with the EZ-96 DNA methylation kit (shallow; Zymo Research Corporation, Irvine, USA). DNA methylation levels were then measured on two arrays: the Illumina Infinium HumanMethylation450 and EPIC BeadChip array (Illumina Inc., San Diego, USA). Preprocessing in ALSPAC was performed with the meffil package.□ Quality control checks included mismatched genotypes, mismatched sex, incorrect relatedness, low concordance with other time points, extreme dye bias, and poor probe detection.

We derived a comprehensive list of methylation age measures in late adolescence.

Specifically, we obtained state measures of methylation age based on:

- Horvath et al. (2013): MethAge_Horvath_,
- Zhang et al. (2019): MethAge_Zhang_,
- Shireby et al. (2020): MethAge_Cortical_ and
- Belsky et al. (2022): MethAge_PACE_.

For each state measure, we derived a predicted age *difference* (PAD) score, which equalled methylation age minus chronological age. We also derived a predicted age *residual* (PAR) score, commonly reported and obtained by regressing chronological age on methylation age. PAR scores can be interpreted as the biological age of a person, which is independent of chronological age. Because our analyses were based on participants younger than 18 years, we did not derive methylation age measures, which were trained exclusively on adult samples (Hannum et al., 2013; Levine, 2013). We also created a Pace of Ageing Methylation (PACE; Belsky et al., 2022) score, which is a rate-rather than a state measure of methylation age. In total, we created seven age measures. For an overview of measures, see Table 2.

**Table 2.**
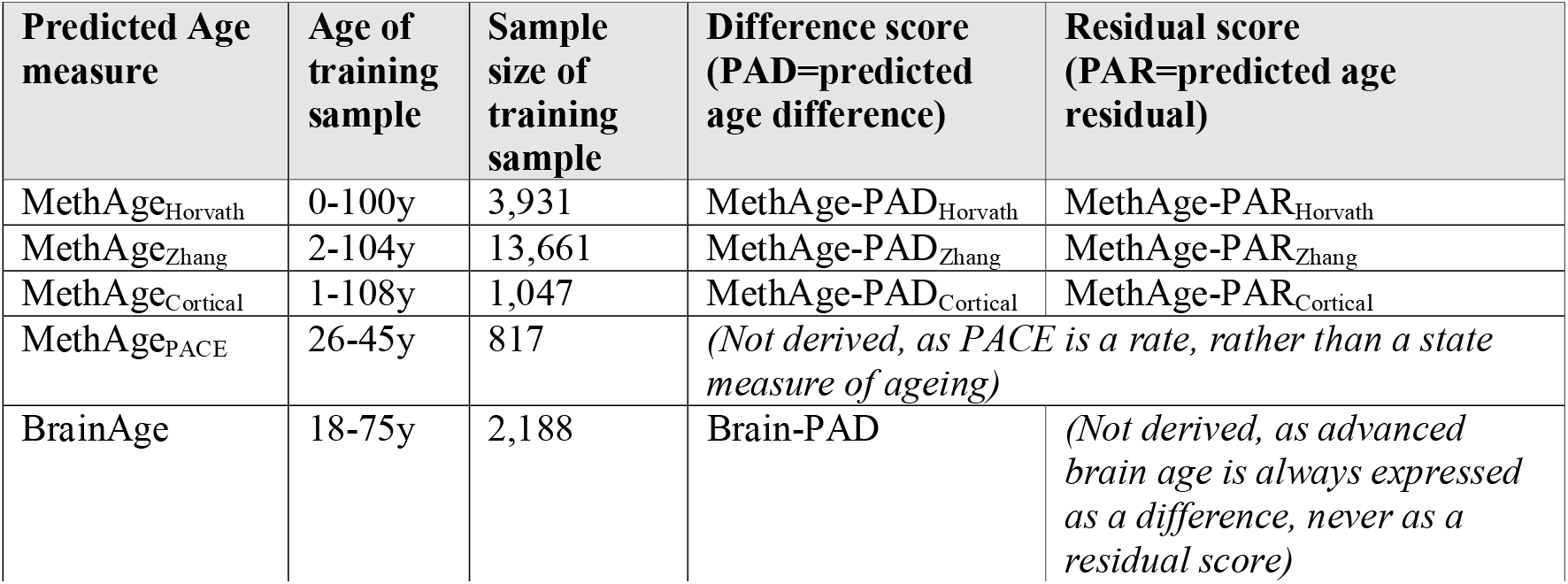
Overview of measures of biological age, assessed in the current study.

| Predicted Age measure | Age of training sample | Sample size of training sample | Difference score (PAD=predicted age difference) | Residual score (PAR=predicted age residual) |
| --- | --- | --- | --- | --- |
| MethAge<sub>Horvath</sub> | 0-100y | 3,931 | MethAge-PAD<sub>Horvath</sub> | MethAge-PAR<sub>Horvath</sub> |
| MethAge<sub>Zhang</sub> | 2-104y | 13,661 | MethAge-PAD<sub>Zhang</sub> | MethAge-PAR<sub>Zhang</sub> |
| MethAge<sub>Cortical</sub> | 1-108y | 1,047 | MethAge-PAD<sub>Cortical</sub> | MethAge-PAR<sub>Cortical</sub> |
| MethAge<sub>PACE</sub> | 26-45y | 817 | *(Not derived, as PACE is a rate, rather than a state measure of ageing)* |  |
| BrainAge | 18-75y | 2,188 | Brain-PAD | *(Not derived, as advanced brain age is always expressed as a difference, never as a residual score)* |

### MRI preprocessing

Between ages 18 to 24 years, a subset of ALSPAC participants were invited to three separate ALSPAC neuroimaging studies (Sharp et al., 2020). In total, MRI data was acquired for n=958 participants, of which n=386 also had methylation data after quality control. For each participant, structural neuroimaging data were acquired on a General Electric 3T HDx scanner. T1-weighted images were processed using the automated FreeSurfer brain imaging software package (Version 6.0.0). Reconstructed images were subjected to quality control measures following the ENIGMA consortium structural image processing protocol (http://enigma.ini.usc.edu/protocols/imaging-protocols/ and Sharp et al., 2020). From these data a measure of brain age was derived in early adulthood (mean=19.8y; SD=1.2y) using ridge regression combining seven subcortical regions, lateral ventricles, and thickness and surface area for 34 bilateral cortical regions (Han et al., 2021). We derived a difference score (Brain-PAD) but not a residual score, as the accepted standard in the field is to describe advanced brain age through a difference score (Franke and Gaser, 2019). In a sensitivity analysis, we validated our results using a different brain age measure (CentileBrain; Yu et al., 2023).

### Health phenotypes

Biological age has been linked to both physical, cognitive and mental health outcomes in adulthood and old age (Bressler et al., 2020; Han et al., 2021; Jansen et al., 2021). To assess these relationships in adolescence, we selected five key measures: (1) C-reactive protein levels as a measure of inflammation (CRP, in mg/L measured at age 17y); (2) BMI (in kg/m^2^ measured at age 17y); (3) smoking behaviour; (4) cognitive performance (WISC performance test score, only available at 8 years); and (5) self-reported depressive symptoms (based on CIS-R measured at 17.5y). Due to the low number of participants who self-reported on their smoking (22%), we used a measure of epigenetically predicted smoking (based on DNAm_cg05575921_), a well-validated proxy for smoking (Bojeson et al., 2017).

### Statistical analysis

Figure 1 displays our analysis flowchart. For Aim 1 and 2, we first assessed the mean deviation and mean absolute error (MAE) between chronological and biological age. Larger deviation or MAE values indicate a larger disparity between chronological and biological age. We then used Pearson correlations to assess the association between chronological age, biological ages, their difference and residual scores. To assess the robustness of these correlations against extreme values, we also applied Spearman rank correlations. We then used principal component analysis to characterise any potential clustering of participants based on their biological age measures, and linear regression to assess the relationships between biological age measures in four regression models with different covariate adjustments:

**Figure 1.**
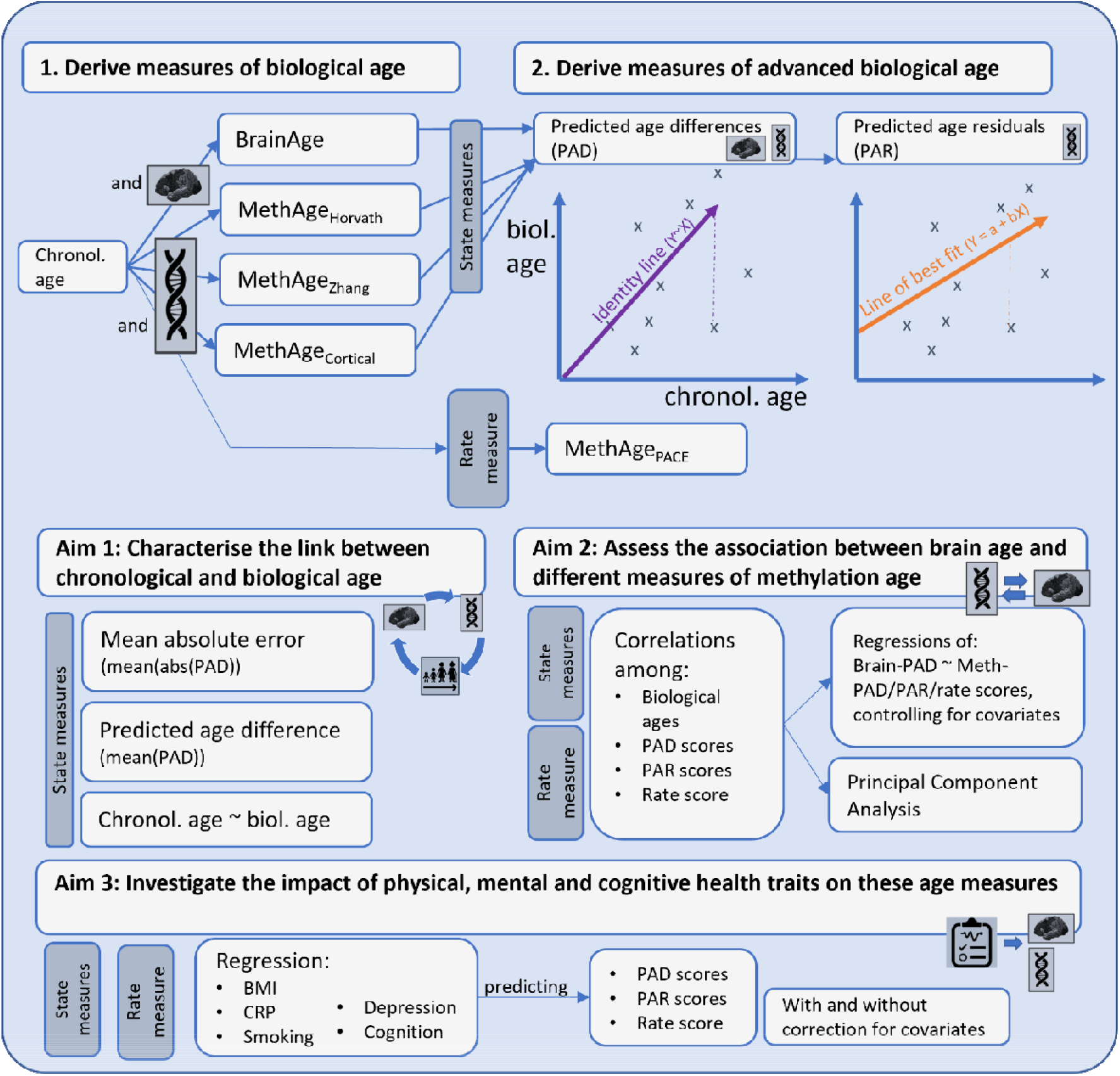
Analysis flow chart.

- Model 1: no adjustment;
- Model 2: controlling for possible batch effects;
- Model 3: controlling for array (450k versus EPIC), MRI study, and cell type (based on Reinius et al. (2012); and
- Model 4: model 3 + age.

Model 4 was carried forward to investigate the relationship between biological ages and health phenotypes (Aim 3). As health traits were obtained either at the same time or earlier than the biological data, health traits were modelled as predictors of biological age. To minimise the influence of extreme outliers, all measures of biological age and health were winsorized based on a cut-off of 5 times the interquartile range. Effectively, this replaced one value for MethAge-PAR_Zhang_ and 17 values for CRP (4.6%).

## Results

### Biological ages capture chronological age to varying degrees in late adolescence

The accuracy of four biological age predictors was acceptable with a MAE between 2.2 and 5.9 years (range: 0.0 to 24.5 years; Figure 2A). The accuracy of MethAge-PAD_Cortical_ was poor (MAE mean=10.2 years; see also SM Table 1). Predictions were all slightly biased towards overestimating ages (Figure 2B and SM Table 1). The smallest average overestimation was observed for MethAge-PAD_Zhang_ (0.9y), followed by MethAge-PAD_Horvath_ (3.7y), Brain-PAD (4.3y) and MethAge-PAD_Cortical_ (10.1y). That is, even though the MAE for most predictors were acceptable, age predictions were generally ‘older’ than chronological age.

**Figure 2.**
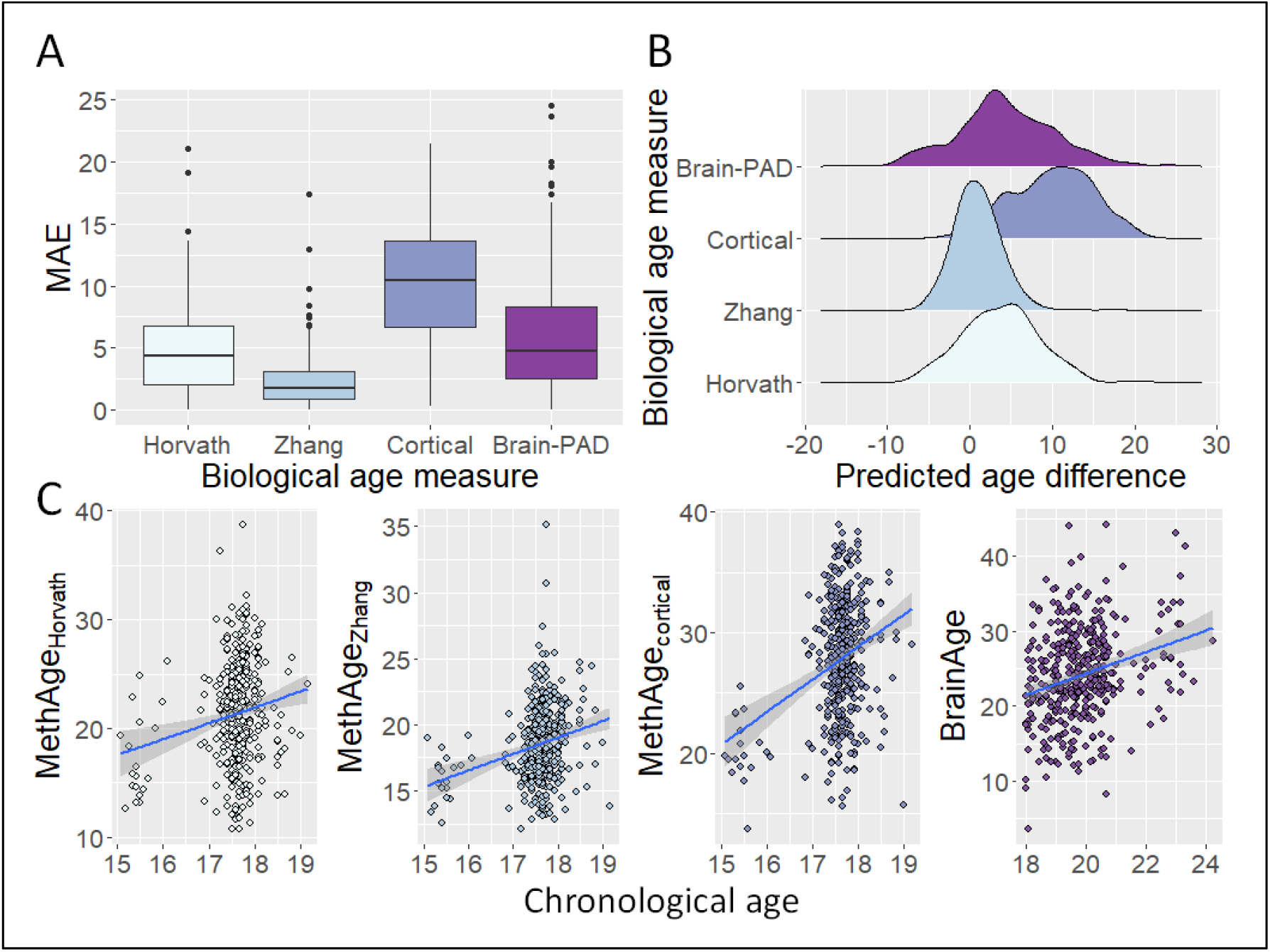
A) Mean absolute errors, B) predicted age differences and C) age correlations across four state measure of biological age.

Predicted ages correlated positively, but not strongly with chronological age, which is unsurprising given the narrow age range of this sample (Figure 2C). The strongest Pearson correlations were observed for MethAge_Cortical_ (r=0.31), followed by BrainAge (r=0.27), MethAge_Zhang_ (r=0.25), and MethAge_Horvath_ (r=0.18). These results were robust to outliers, as similar patterns were observed using Spearman correlations (SM Table 2).

As expected, biological age *residuals* were independent of chronological age (r range: -0.03 to 0.01). However, biological age *difference* scores were also largely unrelated to chronological age (r range: 0.05 to 0.08), except for MethAge-PAD_Cortical_ (r=0.20). Because difference scores correlated strongly with residual scores (all r’s > 0.97), we focused on difference scores (and MethAge_PACE_), while presenting results for residual scores throughout the supplemental materials.

### Brain age is largely independent of methylation ages in late adolescence

Although some biological age measures or their difference scores were correlated with each other (range r: -0.13 to 0.24; SM Table 2 and 3), most associations decreased substantially once controlling for batch, cell type, array or age (and MRI study when BrainAge-PAD was the outcome; SM Table 4). The exception was Horvath’s methylation age and age difference score, which were associated positively with all measures of cortical methylation age (SM Table 4).

These findings suggest that most measures of methylation or brain age are largely independent of each other in late adolescence.

Interestingly, despite being trained on cortical tissue, all measures of cortical methylation age were more strongly associated with other methylation measures in blood tissues (r between -0.13 and 0.24) than with measures of brain age (r between -0.05 and 0.06). This could suggest that biological modality of data collection (methylation versus neuroimaging) might account for more heterogeneity than the tissue of interest (blood versus brain).

BrainAge-PAD remained largely independent of the other methylation PAD scores (Figure 3). Although overall largely insignificant, most associations between BrainAge-PAD and state-based methylation age measures were negative, while the association with MethAge_PACE_, a rate-based measure, was positive. Principal component analyses indicated no clustering of participants based on their biological ages or difference scores (SM Figure 1A-C), but 82% of participants showed consistent accelerated (or decelerated) biological age in more than one measure (SM Figure 1D).

**Figure 3.**
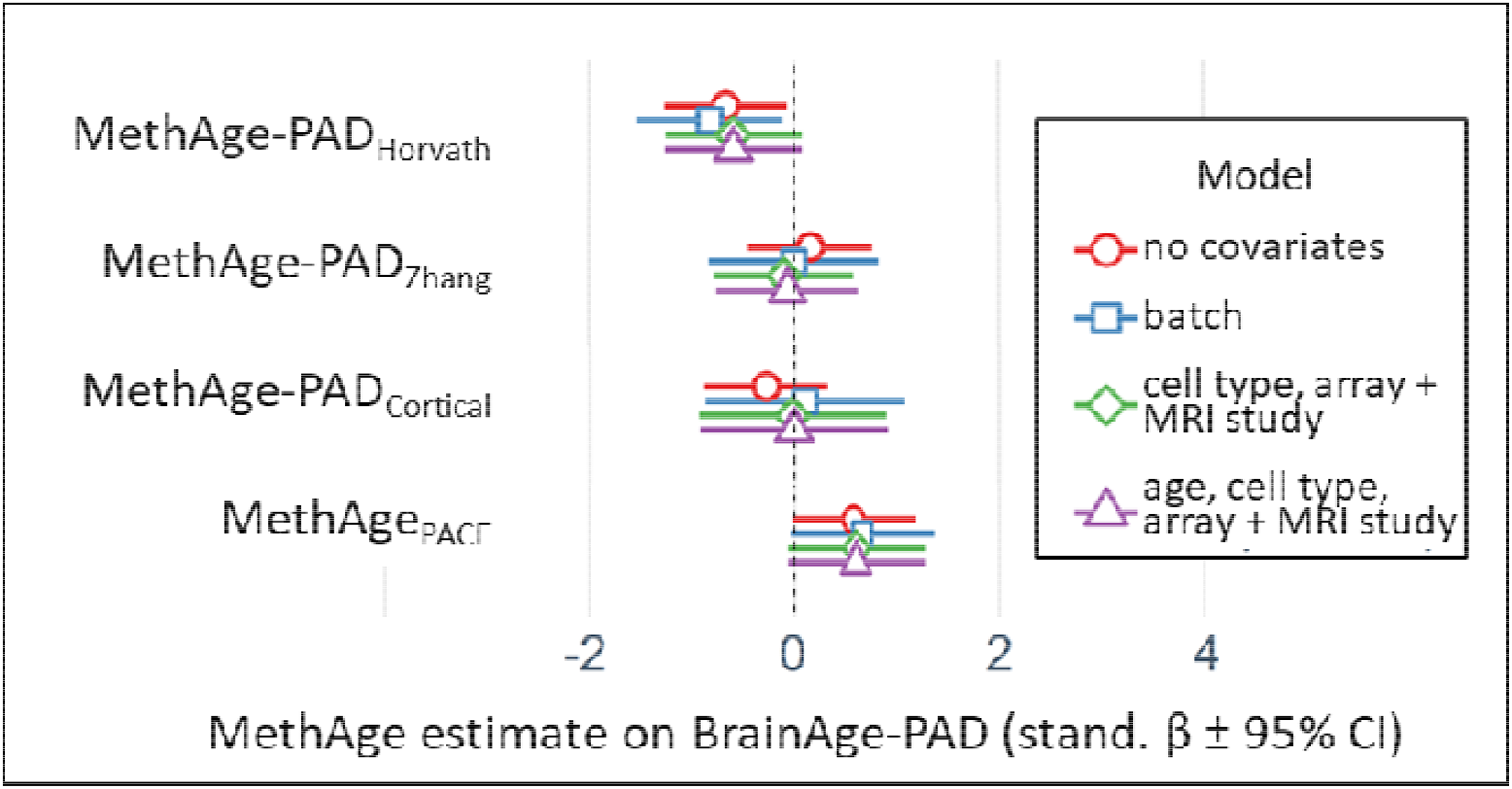
Standardized regression betas (95% CI) of four methylation age difference scores predicting BrainAge-PAD.

### Smoking behaviour and BMI associated with blood-but not brain-related measures of age

A faster rate of ageing (MethAge_PACE_) and advanced methylation age (MethAge-PAD_Zhang_) were linked to increased smoking (beta_std_ between -0.01 and -0.35; Figure 4A-B and SM Table 6), even after controlling for age, array, MRI study and cell type. MethAge-PAD_Horvath_ – and -PAD_Zhang_ were positively associated with BMI (beta_std_ = 0.52; 95% [0.08; 0.95] and beta_std_ = 0.27; 95% [0.01;0.52]; Figure 4C-D and SM Table 6).

**Figure 4.**
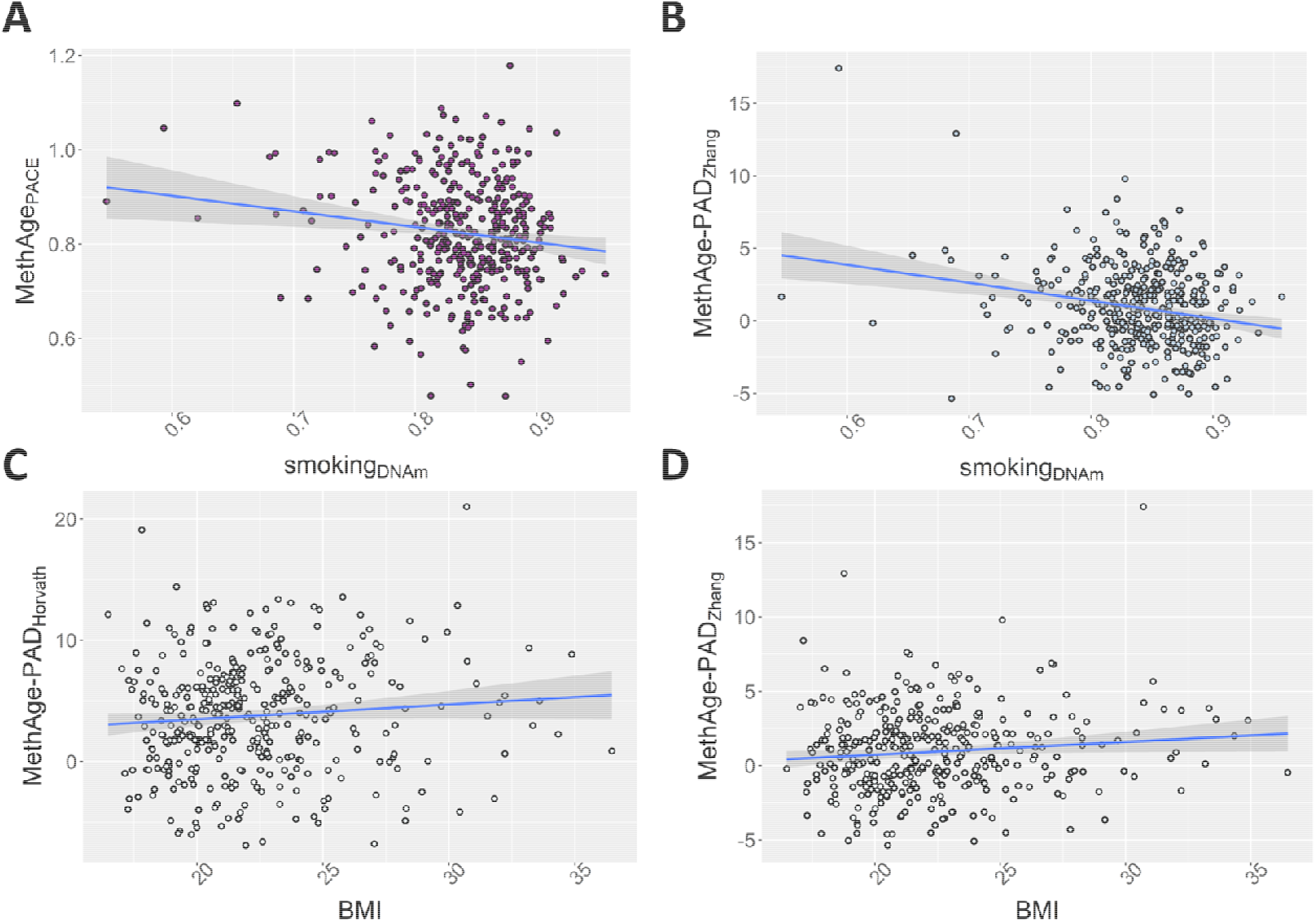
A-B) Smoking behaviour and C-D) BMI associated with three measures of methylation age. Due to the low number of participants who self-reported on their smoking (22%), we used a measure of epigenetically predicted smoking (smoking_DNAm_, based on DNAm_cg05575921_), a well-validated proxy for smoking. Lower smoking_DNAm_ values associate with increased smoking behaviour.

All remaining associations between biological ages (including all difference or residual scores of methylation or brain age) and measures of physical (CRP), cognitive or mental health (cognition, depression) were weak and insignificant (standardized regression betas range between -0.37 and 0.43; SM Table 6).

A similar pattern of results was obtained when using the Centile measure of BrainAge-PAD (SM Figure 2 and SM Tables 7-8).

## Discussion

In this study, we comprehensively characterised the link between methylation age and brain age in adolescence. We highlight three key findings: 1) multiple measures of biological age captured chronological age in late adolescence; 2) there was little evidence that measures of methylation age and brain age were related and 3) smoking and BMI predicted advanced methylation age, but not brain age.

First, chronological age was associated with biological age (including both methylation and brain age) to varying degrees in late adolescence. With an age correlation between 0.18 to 0.31 and an MAE between 2.2 and 10.2 years, overall biological age measures were moderate-to-good indicators of chronological age in late adolescence. We note, however, that on average biological age predictors overestimated chronological age by 1 to 10 years. Although large, the range of errors are consistent with previous research (Hannum et al., 2013; Horvath et al., 2018; Marioni et al., 2019). Sample differences may explain some of these large MAEs. For example, the largest MAE was observed for MethAge_Cortical_, which was developed in cortical tissues on a smaller training sample and had lower accuracy when applied to non-cortical tissues such as blood (Shireby et al., 2020). Conversely, the lowest MAE was observed for MethAge_Zhang_, which was developed in a large training sample and based on blood and saliva tissue. These results further suggest that tissue and training sample size are important factors affecting prediction accuracy. Furthermore, the median age in the training sample for MethAge_Cortical_ was 57 years – over 15 years older than the median age of the training samples used for MethAge_Horvath_.

Despite these caveats, we observed consistency across measures. Most people, who showed advanced age in one measure, also did so in other measures. Furthermore, as argued elsewhere (de Lange et al., 2022), performance metrics such as MAE are sample-specific (i.e., they cannot easily be compared across studies) and should be considered in combination with other metrics such as correlation coefficients. For example and similar to our study, large increases in MAE have been reported despite acceptable correlations (r > 0.5) between chronological and predicted age, especially in situations where the age range of the testing sample was very narrow and different to that of the training sample (de Lange et al., 2022). In line with this, we observed that – even though MethAge_Cortical_ had the largest MAE across all biological ages – it was also the measure most strongly associated with chronological age and Horvath’s methylation age. Hence, a large MAE should not be considered the only indicator for model accuracy.

The degree to which age deviations are influenced by genetic or environmental factors remains unclear. Previous research reported decelerated methylation age clustering within families (Horvath et al., 2015) and a partial genetic basis underlying some, but not all measures of methylation age (Gibson et al., 2019). Other research highlighted the impact of environmental factors such as childhood adversity in predicting advanced methylation age (Marioni et al., 2019). Therefore, the large, predicted age differences reported in our study perhaps indicate the extent to which biological age can diverge from chronological age already in late adolescence, either because of genetic factors, environmental variation in childhood experiences, or due to stochastic effects, which should be investigated in future research. This emphasises the importance of understanding the ageing process during the first two decades of life to create interventions that can reduce ageing-related health inequalities seen in later life (Deckers et al., 2019).

The second key finding highlights that brain age was found to be largely independent of methylation ages in late adolescence. This interpretation is consistent with Teeuw et al. (2021), Zheng et al. (2022) and Cole et al. (2018), whereby only weak associations were detected between brain age and methylation age in a sample of younger or older adults. Our findings, however, do contradict previous research in older individuals, which reported associations between neuron density or brain vascular lesions, as measures of brain age, and methylation age (Hillary et al., 2021; Lu et al., 2017).

Research into the blood-brain barrier may provide an insight into some of these inconsistencies. The blood-brain barrier is most intact in early development and only degrades in later life, resulting in increased permeability of the blood-brain barrier with age (Farrall and Wardlaw, 2009). This increased permeability could affect neuron density (Sweeney et al., 2018), the cellular measure of brain age used in Lu et al. (2017), but might relate less strongly to the brain imaging measure of brain age used in the current study. Hence, it is possible that methylation age associates with microscopic (cellular), rather than macroscopic measures of brain age, especially in later life, when the blood-brain barrier might be less intact (Farrall and Wardlaw, 2009).

It is also possible that differences in findings are accounted for by the use of diverging measures of methylation age. Hillary et al. (2021) used *GrimAge*, a methylation age measure predictive of mortality, rather than chronological age. We did not use this measure, as it was trained on adults only and hence did not apply to the age range of our adolescent sample.

Consequently, more research is recommended to test the independence between methylation age and brain age using a range of different cell-based and neuroimaging measures.

The third key finding indicated that advanced biological age in late adolescence is only weakly linked to health traits. Of all tested traits, smoking behaviour was most consistently associated with measures of methylation age, including the pace of methylation ageing, but not with brain age. The association with methylation age is consistent with previous research, as smoking has been associated with advanced methylation age, specifically in lung tissue (Wu et al., 2019). The lack of association between smoking and brain age however appears to be inconsistent with previous findings, whereby brain volume loss or brain age has been reliably associated with smoking in middle-aged adults (Durazzo et al., 2017; Elbejjani et al., 2019; Ning et al., 2020). It is possible that effect of environmental exposures such as smoking on the brain do not become apparent until later in life. Additionally, duration of exposure could be relevant, as adolescents will have smoked on average for a shorter period than adults, limiting the time for smoking to reliably affect tissues more distal from the lungs (Szulc et al., 2002). Finally, methodological considerations may also contribute to the lack of association between smoking and brain age. Although we used a validated biomarker of smoking, this measure is derived from methylation data. This may lead to a stronger association of smoking with methylation age as opposed to brain age.

Our study should be considered in light of the following limitations: the age range of our sample was very narrow, making it difficult to disentangle measurement error from true predicted age deviations. However, all measures of biological age were associated with chronological age despite the narrow age range. This suggests that the non-association across biological ages, we observed, is likely a true finding. Second, our sample was limited to the period of late adolescence. It is possible that biological age measures associate differently with each other or health traits at different developmental periods. However, using a sample with a highly constrained age range allowed us to derive developmentally specific conclusions. It might also have minimised some of the noise that could have obscured an association in studies with a larger age range. Third, the sample size – although similar to previous studies – was moderate. Future research is needed to replicate our results in larger samples of individuals across a wider age range.

In summary, we investigated the association between chronological age and two modalities of biological age - measures of methylation and brain age in adolescents. The variability of and independence between these age measures in adolescents highlights the dynamicity of ageing across the lifespan and signifies the importance of tracking the mosaic of ageing already in younger populations to inform prevention targets to support physical and mental health across the life course.

## Acknowledgements

We are extremely grateful to all the families who took part in this study, the midwives for their help in recruiting them, and the whole ALSPAC team, which includes interviewers, computer and laboratory technicians, clerical workers, research scientists, volunteers, managers, receptionists and nurses.

